# Characterizing and correcting camera noises in back-illuminated sCMOS cameras

**DOI:** 10.1101/2021.01.01.425025

**Authors:** Zhaoning Zhang, Yujie Wang, Rafael Piestun, Zhen-Li Huang

## Abstract

With promising properties of fast imaging speed, large field-of-view, relative low cost and many others, back-illuminated sCMOS cameras have been receiving intensive attentions for low-light imaging in the past several years. However, due to the pixel-to-pixel difference of camera noises (called noise non-uniformity) in sCMOS cameras, researchers may hesitate to use them in some application fields, and sometimes wonder whether they should optimize the noise non-uniformity of their sCMOS cameras before using them in a specific application scenario. In this paper, we systematically characterize the impact of different types of sCMOS noises on image quality and perform corrections to these sCMOS noises. We verify that it is possible to use appropriate correction methods to push the non-uniformity of major camera noises, including readout noise, offset, and photon response, to a satisfactory level for conventional microscopy and single molecule localization microscopy. We further find out that, after these corrections, global read noise becomes a major concern that limits the imaging performance of back-illuminated sCMOS cameras. We believe this study provides new insights into the understanding of camera noises in back-illuminated sCMOS cameras, and also provides useful information for future development of this promising camera technology.

## 1. Introduction

Low-light cameras are indispensable for various low-light imaging applications, especially single molecule fluorescence microscopy [1]. Semiconductor complementary metal oxide semiconductors (sCMOS) camera is a new type of low-light cameras with high imaging speed and large field-of-view, and thus is well-suited to be used in high-speed fluorescence microscopy of biological samples.

However, the detectability of sCMOS cameras is limited by their relatively low signal-noise-ratio (SNR), especially in comparison with another popular type of low-light cameras: electron multiplier charge-coupled device (EMCCD) cameras [2]. It is well-known that SNR could be improved by either increasing quantum efficiency (QE) or decreasing camera noises. Recently, with the invention and mature of back-illuminated sCMOS sensors, the QE of sCMOS cameras has been increased to ∼ 95%, and the SNR of some commercial back-illuminated sCMOS cameras is even better than that of many EMCCD cameras when the incident signal is > 4 photon/pixel [3]. Clearly, decreasing camera noises becomes the next step for further pushing the detectability of back-illuminated sCMOS cameras into single-photon detection regime.

Compared with EMCCD cameras, sCMOS cameras suffer from not only higher global readout noise, but also larger pixel-to-pixel difference in camera noises (that is, noise non-uniformity). The latter is mainly originated from the individual readout structure in sCMOS cameras [4]. Various efforts were developed to characterize and correct the noise non-uniformity. The camera noises in sCMOS cameras (abbreviated as sCMOS noises hereafter) were characterized [5–8] and their impacts on some applications, such as single molecule localization microscopy (SMLM), were studied [9–14]. Noise correction algorithms were integrated into some commercial sCMOS cameras by their manufactories, or developed for several specific imaging scenarios by researchers [11, 13, 15–17].

Since the noise correction algorithms developed by researchers were usually designed to correct several types of sCMOS noises simultaneously, it is difficult to obtain quantitative information on what level a specific type of sCMOS noises has been corrected to. Besides, some noise correction algorithms (for example, defect pixel correction) have been integrated into commercial sCMOS cameras, and are routinely used by many users without any precautions. However, it is not clear whether these noise correction algorithms should be used in some special application scenarios. Moreover, it is generally believed that the characteristic of sCMOS noises under long exposure time is different from that under short exposure time, but the performance of the noise correction algorithms under different exposure times has not been well studied. In a word, although many noise correction algorithms have been developed and used to improve the noise non-uniformity of sCMOS cameras, researchers are still confused about when and how to select appropriate noise correction algorithms in their specific applications. They may even have no confidence on which technical specifications in a camera datasheet are more important for choosing an appropriate camera, and whether the noise correction algorithms used by their colleagues should be modified before being used in their specific experiments.

In this paper, we systematically analyze the impact of different types of camera noises on image quality, and evaluate whether a specific camera noise can be properly corrected. Firstly we characterize individual camera noises in two popular back-illuminated sCMOS cameras and a popular EMCCD camera. Then, we investigate the impact of different types of camera noises on SMLM and conventional microscopy. We take special efforts on analyzing the global read noise and read noise non-uniformity. Finally, we quantify some commonly-used noise correction algorithms. We confirm that the impact of noise non-uniformity on image quality could be minimized to a negligible level. After applying these noise corrections, we find out that global read noise becomes the major camera noise that limits the imaging performance of back-illuminated sCMOS cameras.

## 2. Theory and Methods

### 2.1 The background of sCMOS noises

There are mainly three types of noises in an sCMOS camera: fixed pattern noise (FPN), read noise, and shot noise [9]. FPN represents the pixel-to-pixel difference of time-independent fixed bias, and can be further divided into offset FPN and gain FPN, which account for pixel-dependent variations in dark signal and photon response, respectively [18]. Read noise usually represents all of the camera noises that are independent on signal intensity [19]. To distinguish read noise from offset FPN, here we only consider the signal-independent *temporal* noise as read noise. Shot noise is originated from quantum fluctuation, and is always equal to the square root of input signal. Since shot noise is from incident light itself instead of the associated camera, we would not consider it as camera noise in this work.

In an sCMOS camera, light hitting the sensor is converted into photoelectrons, and then into voltage by the individual voltage converter in each pixel. Moreover, each column has its own Analog to Digital Converter (ADC). This kind of readout structure increases the pixel-to-pixel difference of offset, photon response and read noise, and thus resulting in a more severe noise non-uniformity. Camera manufactories usually provide dark-signal non-uniformity (DSNU), photon response non-uniformity (PRNU), and root-mean-square (RMS) value of read noise to characterize the amplitude of offset FPN, gain FPN, and read noise, respectively. However, the RMS of read noise can only characterize the amplitude of global read noise, and there is no parameters to characterize the read noise amplitude variation from pixel to pixel. In this work, to keep consistent with DSNU and PRNU, we propose to use read noise non-uniformity (RNNU), which is calculated by the ratio of the standard deviation (SD) to the RMS of read noise, to describe the read noise amplitude variation from pixel to pixel. We compare the impact of global read noise and RNNU on imaging quality.

For pixel *i* in an sCMOS camera, when the incident signal is *S*_*i*_, the output digital value is modelled as:

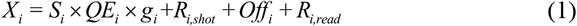

where *R*_*i,shot*_ denotes shot noise and *R*_*i,read*_ denotes read noise, and both of them are temporal noise; *g*_*i*_ and *Off*_*i*_ are gain and offset value, respectively. Note the difference of *QE*_*i*_, *g*_*i*_, *Off*_*i*_, *R*_*i,read*_ from pixel to pixel are used to characterize the noise non-uniformity in the sCMOS camera. Dark current is not included in this model because: 1) most of the experiments in this work are performed with short exposure time (≤ 20 ms), where dark current is negligible; 2) for any given exposure times, the major impact of dark current on output digital value is already included in the amplitude of offset and read noise.

Defect pixels are pixels with abnormal performance that disturb users or even arouse imaging errors. However, the standard for distinguishing abnormal from normal performance is usually not clear. Generally, there are two types of defect pixels: high dark noise pixels and low gain pixels [20, 21]. High dark noise pixels are pixels with high dark electric signal or dark noise variation, including but not limit to high dark current pixels (also called hot pixels). Low gain pixels are pixels with relatively low photoelectric conversion ability. Clearly, defect pixels result in high noise non-uniformity. Note that the number of defect pixels usually increases during the manufacturing process or during the usage of an sCMOS camera [20, 21]. Besides, since defect pixels (or high noise pixels) are easily observed in enlarge images, they were often used to analyze the impact of noise non-uniformity on imaging quality [11, 13, 16, 17].

### 2.2 Theoretical background for sCMOS noise characterization

We use three noise maps (including photon response map, read noise map and relative offset map) to characterize the sCMOS noises in every pixel. Based on Eq (1), when there is no incident light (called dark frame), the output digital value is only determined by the offset and read noise. For each pixel, we use the mean and the SD values of continuous *N* dark frames to represent the offset value (*Off*_*i*_) and read noise value (*σ*_*i,read*_) of a pixel, respectively:

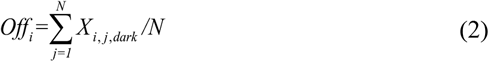

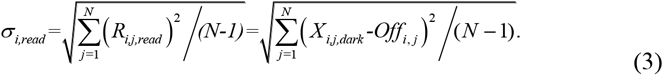

*Off*_*i*_ and *σ*_*i,read*_ can be further transferred to electron unit by dividing their grey values with the camera gain value, and the latter is usually provided by camera manufactory or measured by PTC method [5]. To help compare the offset values between different cameras, we use relative offset, that is, the offset subtracted with the mean offset value of all pixels. The number of dark frames used to calculate the read noise and offset is 5000 for normal exposure time (1s or shorter) and 1000 for long exposure time (> 1s).

We use a uniform-illumination system to measure the photon response map. We assume that the photon signal is the same for all pixels in a frame (*S*_*i*_ = *S*), and use the averaged digital value 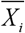 from continuous frames to eliminate shot noise and read noise. We calculate the relative photon response value (*rp*_*i*_) of each pixel as the ratio between the signal value of a single pixel to the mean value of all the pixels:

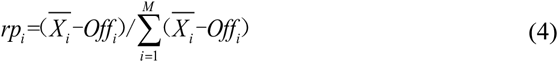

where *M* is the number of all pixels. To calculate the relative photon response from several groups of images captured at different signal intensity levels, we use a linear fitting with 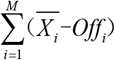 as dependent variable and 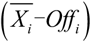 as independent variable for each pixel. Usually we capture 6 groups of raw images with 1000 frames in each group. The illumination intensities are usually controlled to uniformly distribute among 10% ∼ 85% of the full signal range. If necessary, the illumination intensities can also be adjusted to provide a photon response map that matches with experimental conditions.

We calculate the precision of noise map measurement by [5]:

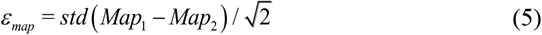

where *Map*_*1*_ and *Map*_*2*_ refer to two maps measured independently under the same experimental conditions, and *std* is the SD of all pixels. The two maps are recommended to be measured at different days to account for experimental environment changes.

To assess the noise non-uniformity of sCMOS cameras, we capture continuously a large number of dark frames to calculate relative offset map, and bright frames at one signal level (typically, half of the signal range) for photon response map. Under the non-uniformity nomenclature, the relative offset map and photon response map here are referred as DSNU map and PRNU map, respectively.

DSNU and PRNU are two widely-used parameters for quantitative camera assessment, as mentioned in the camera calibration standard EMVA 1288 [22]. DSNU is defined as the SD of dark signal and can be calculated by the SD of the relative offset map (DSNU map) in electron unit:

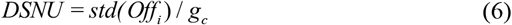

where *g*_*c*_ is the camera gain value, and *std* is the SD of all the pixels. PRNU is defined as:

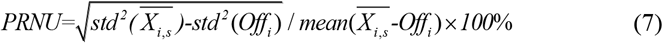

where 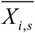 represents the mean digital value of pixel *i* measured at signal intensity level *S*, and *std* is the SD of all pixels. PRNU equals to the SD of PRNU map (or photon response map measured at one signal intensity level *S*). In this work, to assess the PRNU of a camera at full signal range, we also calculate the SD of the photon response map measured from several different signal levels.

We also use local sensitivity variation (SV) to characterize local photon response non-uniformity [13], and calculate it as:

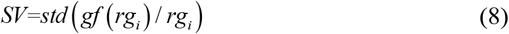

where *gf* represents Gaussian smoothing filtering with a sigma of 9 pixels, and *std* is the SD of all pixels. Compared with PRNU, SV only characterizes the local photon response non-uniformity, and thus is suitable for applications with a small number of pixels, such as SMLM.

We calculate RNNU as:

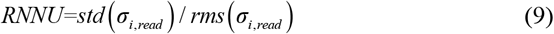

where *std* and *rms* are the SD and RMS of all pixels, respectively.

### 2.3 Experimental conditions for sCMOS noise characterization

A customized system based on an integrating sphere was specially designed for characterizing the PRNU of low-light cameras. Three LEDs with peak wavelength around 400 nm (EP-U4545K-A3, Epileds, China), 600 nm (BN-R3838C-A3, Epileds, China), and 850 nm (ES-SASFPN35, Epileds, China), respectively, were placed at the input port of an integrating sphere (Flight technology, China) as the light source. The LED intensity was controlled by home-built electronic circuits. To decrease the irradiance difference between the camera edge and the camera center, the tested low-light cameras were placed ∼30 cm away from the exit port (diameter: ∼8 cm) of the integrating sphere. The camera sensor should be aligned to be completely vertical to a plane where the longitudinal axis of the exit port lies; however, we found it is hard to obtain such a perfect alignment, and the residual angle would degrade the effectiveness of PRNU correction. Therefore, we fixed the test camera on the platform for several days to guarantee the same illumination angle for PRNU map and photon response map measurement. The input port (from the LEDs to the integrating sphere) and the exit port (from the integrating sphere to the test camera) of the integrating sphere were covered separately to block ambient light, so that the tested camera received photons only from the LEDs. The threaded metal adapter on the low-light cameras was removed to minimize illumination uniformity deterioration. The time fluctuation of the illumination was measured to be 0.05% root-mean-square (RMS) using a Flash 4.0 V3 (SN: 303487, Hamamatsu Photonics, Japan) at the intensity of ∼15000 photons/pixel.

Two popular back-illuminated sCMOS cameras were tested in their best modes for low-light detection: the high gain mode for a Dhyana 95 (SN: KBS4951703002, Tucsen Photonics, China), and the 12 bit high sensitivity mode for a Prime 95B (SN: A18A203022, Photometrics, USA). The camera gains for these two modes, measured from the PTC method [5], were 1.98 DN/e-for the Dhyana 95 and 1.64 DN/e-for the Prime 95B, respectively. The maximum signal range for these two modes are ∼ 2000 e-/pixel for the Dhyana 95 and ∼ 2400 e-/pixel for the Prime 95B. The exposure times were all set to be 20 ms except otherwise specified.

### 2.4 sCMOS noise correction

In addition to the normal denoising tasks, an sCMOS denoising algorithm should also consider noise non-uniformity. Note that FPN correction and defect pixel correction are usually performed by CMOS camera manufactories to satisfy their users. For both the FPN and defect pixel correction algorithms, we classify them into two types: static correction algorithm and dynamic correction algorithm. The only difference between them is on how to calibrate the camera noises. In the static correction algorithm, special images (typically under homogeneous illumination [20, 23]) are taken and used to calculate the characteristics of the camera noises. In the dynamic correction algorithm, the camera noises are characterized from normally images [20, 21, 23, 24]. Then, different noise correction strategies are used according to the camera noise characteristics. The static correction algorithm is usually used in sCMOS camera, after considering the following reasons: 1) sCMOS cameras have a more stable noise characteristic than normal CMOS cameras, thus the static noise correction in sCMOS cameras can be valid for a long time; 2) Raw data is desirable in many applications using sCMOS cameras, but the dynamic correction algorithm usually processes more raw data than the static correction algorithm.

In this paper we adopt two common-used static correction algorithms: a modified two-point correction algorithm for FPN correction, and a static local mean filtering algorithm for defect pixel correction. The normal two-point correction algorithm assumes a stable and linear photon response from each pixel [25]. It first captures images under homogeneous illumination and two representative signal levels, then uses them to calculate the offset map and the photon response map. In the modified version, the offset map is calculated from dark images, and the photon response map is calculated from bright images at one or several signal levels [13, 14].

These two maps are further used to correct the raw images via an inverse operation of Eq. (1), and can be expressed as:

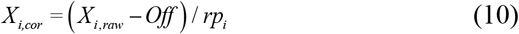

where *X*_*i,cor*_ and *X*_*i,raw*_ are the corrected digital value and the raw digital value for pixel *i*, respectively. When there is no photon response map, Eq. (10) can be simplified as:

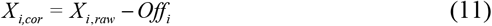

For defect pixel correction, a common strategy used in both dynamic and static correction algorithms is that the raw digital value of a defect pixel is substituted by a mean value calculated from the surrounding pixels [21, 26]. In this work, we first determine the defect pixels from their statistical noise characteristics, and then substitute their values by the mean values of the surrounding 8 pixels. This correction algorithm is thus called static local mean filtering. We consider a pixel as defect pixel if the amplitude of any camera noises in this pixel exceeds 10% of the maximum signal range of the working camera mode. More specifically, for high gain mode in Dhyana 95, defect pixels include: 1) high read noise pixels and high relative offset pixels where the amplitude is > 200 e-under 10 s exposure time, and 2) low gain pixels where the relative photon response is < 0.8 under 20 ms exposure time. Because both read noise and offset increase with exposure time, here the high read noise pixels and the high relative offset pixels are both measured with 10 s exposure time, which is the maximum exposure time of both sCMOS cameras for normal users. However, an exposure time of 20 ms is used for characterizing the low gain pixels for the following reasons: 1) the relative photon response changes only slightly with the exposure time; 2) the offset value is small under short exposure time, leaving sufficient capability for the pixel to response to the incident photons. In addition, the threshold of the relative photon response is set to be 0.8, because a pixel with photon response < 0.8 would lead to a bias that exceeds 10% of the maximum signal range, while the relative photon response is usually measured at half signal range.

Although FPN and defect pixels are usually corrected by sCMOS camera manufactories, researchers may perform FPN re-correction or develop some sCMOS specific algorithms for conventional microscopy and SMLM. Therefore, we evaluated the noise correction ability of two popular sCMOS specific algorithms, including NCS [16] and MLE_sCMOS_ [11]. For conventional microscopy, we chose PURE [27] to compare with NCS. PURE is a denoising algorithm that considers camera noises, but not the noise difference between pixels. Note that NCS uses the noise maps (including offset, read noise, and gain) from every pixel, and combines them with a high frequency filter for denoising, while PURE only uses the noise values (including offset, read noise, and gain) of an entire camera, and combines them with a mixed Poisson-Gaussian model for denoising. For SMLM, a conventional localization algorithm is usually used for calculating the center positions of the molecules in raw images, and shot noise is generally considered in this localization algorithm. The sCMOS specific localization algorithm considers not only shot noise model, but also read noise and FPN. We compared a conventional MLE-based localization algorithm, which is embedded in a widely-used software called ThunderSTORM [28] (referred as MLE_normal_ below), with an sCMOS specific localization algorithm called MLE_sCMOS_ [11].

We evaluated NCS [16] and MLE_sCMOS_ [11] using the Matlab codes provided in the published papers, and tested PURE [27] and ThunderSTORM [28] with the ImageJ plugins provided on the websites. Most parameters were used as the default settings. We replaced the camera noise data with our measurement or simulation. The NCS and MLE_sCMOS_ use a gain map to correct FPN, which is considered to be not accurate enough [3, 15]. Therefore, instead of using the gain map, we used the measured gain value of the camera multiplying by the photon response map.

To compare MLE_normal_ with MLE_sCMOS_, we set the parameters in ThunderSTORM based on those used in MLE_sCMOS_. We chose “Difference of averaging filters” as the image filter, “local maximum” as the localization method, and “Maximum likelihood” as the fitting method. The rendering method was “Normalized Gaussian” for both MLE_normal_ and MLE_sCMOS_. We identified the localized molecules from different algorithms as molecule pairs using the following criteria: two molecules in the same image frame have a distance of less than three pixels (330 nm), but were localized from different localization algorithms. Besides, we adapted the “log-likelihood ratio threshold” in MLE_sCMOS_, which is based on the signal-noise-ratio in the single molecule images. Most parameters were kept the same as the default settings.

### 2.5 Image simulation

Camera noise maps were simulated based on the noise maps of the Dhyana 95 with 20 ms exposure time. To analyze the impact of different camera noises on image quality, the noise maps were scaled to simulate images with different noise amplitudes. To obtain noise maps with different non-uniformity, the SD of the standard maps (the photon response map, read noise map, or relative offset map of the Dhyana 95) was modified by keeping the mean value of the map and scaling the residual of each pixel. To obtain read noise maps with different amplitude of global read noise, the RMS of the standard read noise map was directly scaled. To investigate the impact of different kinds of camera noises on imaging, we changed only one of the three noise maps from the standard map of simulated images. The quantum efficiency (QE) was set to be 82%, which mimics the QE of the Dhyana 95 or Prime 95B around 700 nm. The pixel size was 110 nm.

To simulate conventional microscopy images, we first obtained a ground-truth image from the following steps: average a group of experimental microtubule images, perform FPN correction, and convert the unit from digital number to photon. The mean value of the top 1%∼70% pixels was used as the photon signal value of this ground-truth image. Then, the intensity of the image was scaled to provide images with different signal intensities, and a background photon of 10 photons/pixel was added to the images. Finally, several groups of images with 100 frames in each group were generated based on Eq. (1).

For SMLM, we simulated several groups of single molecule images with different noise maps or signal intensities. Each group contains 400 image frames, and each frame has 400 emitters. The camera noise maps and the ground-truth positions of the emitters were not changed inside the same group. The image size of each frame is 420 × 420 pixels. The camera noise maps were constructed from 400 different areas of 21 × 21 pixels, which were randomly chosen from the scaled noise maps of the Dhyana 95. There was only one emitter in each area (21 × 21 pixels), and the emitter was usually placed randomly in the center pixel (110 nm × 110 nm). For the simulated images with isolated high noise pixels, the center pixel in an area of 21 × 21 pixels was replaced by a high noise pixel in the noise map, and the emitter was set to be located randomly in the area, with a fixed distance from the center of the high noise pixel. Based on the ground-truth positions of the emitters and the signal intensities, a Gaussian model with a sigma of 1.3 pixel was used as the emitter model to present a photon image. A background photon of 10 photons/pixel was then added to the photon image. Finally, simulated image frames were generated using Eq. (1).

### 2.6 Imaging experiments

We performed conventional microscopy imaging on an inverted microscope system (IX-73, Olymplus, Japan) equiped with a 640 nm laser (LWRL640-3W, Laserwave, China), an oil-immersion objective (100×, NA 1.4, Olympus, Japan), and the Dhyana 95 sCMOS camera. The microtubles of U-2 OS cells were labled with Alex Fluor 647 using a typical immunofluorecence method. Fluorescent beads (F8807, FluoSpheres, Molecular Probes, USA) with peak emission wavelength of ∼ 680 nm were fixed on a glass slide. The illumination intensity of the laser was controlled to be low, so that the emission from fluorescent beads is weak and the impact of camera noises on fluorescence images could be clearly visualized. The exposure time was 20 ms for the microtuble imaging and 10 s for the fluorescent bead imaging. The pixel size was 110 nm.

SMLM imaging was performed on the same optical microscope system. Alexa 647 labeled U-2 OS cells were imaged with a standard SMLM buffer as described in our previous work [29]. Raw image frames were captured with 1 ms exposure time to enhance the impact of camera noises on raw images.

### 2.7 Image quality assessment

We use a temporal pixel fluctuation map and three parameters, including peak signal-noise-ratio (PSNR), structural similarity (SSIM), and the number of “outlier pixel” (N_OP_), to assess the quality of conventional microscopy images. PSNR and SSIM are two common-used parameters for comparing de-noising algorithms [30]. PSNR is defined as the ratio between the peak signal value to the noise value:

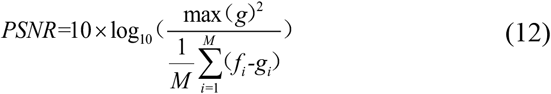

where *M* is the total number of pixels, *f* is the reference image and *g* is the test image. *max(g)* is the peak signal value of the test image. We use the value of the top 1% pixels in the image as the peak signal value to avoid interference from defect pixels. SSIM is calculated as:

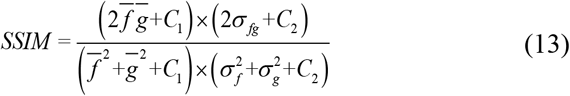

where *σ*_*f*_ and *σ*_*g*_ are the standard deviations and *σ*_*fg*_ is the cross-covariance for the reference image *f* and test image *g*; 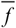and 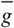 are local mean of the image; C_1_ and C_2_ are used to avoid a null denominator, and are both set to 0.01. For each imaging condition, usually we capture a group of 100 image frames to assess the image quality. We use the mean value of PSNR and SSIM as the figure of merit.

To characterize the temporal noise non-uniformity in conventional microscopy, we calculate the SD value of each pixel in each group of images to present a temporal pixel fluctuation map [16], and use *N*_*OP*_ to characterize the impact of RNNU on conventional microscopy images. Because the pixel values in the temporal pixel fluctuation maps varies gradually, high read noise pixels can be discovered by comparing their pixel values in the temporal pixel fluctuation map with their surrounding pixels. We define “outlier pixel” to be the pixel with a significantly higher value than the surrounding 8 pixels in the temporal pixel fluctuation map:

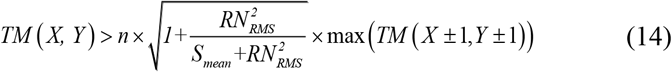

where *TM (X,Y)* is the value of pixel *(X,Y)* in the temporal pixel fluctuation map, *RN*_*RMS*_ is the RMS of read noise, and *S*_*mean*_ is the mean value of the raw image. *n* is an empirical factor to control the dividing line of the outlier pixel, and is set to be 1.25.

To compare the performance of different localization algorithms, we imported the localization results (including but not limit to localization position, background, intensity, uncertainty) of MLE_sCMOS_ into ThunderSTORM, and obtained a rendered super-resolution image. To assess the impact of camera noises on SMLM images, for each experimental conditions, 400 image frames with 400 emitters in each frame were simulated. For each emitter, the SD of the localized positions was calculated as localization precision, and the distance between the mean value of the localized position and the ground truth position was calculated as localization bias. For each group of raw images, we used the RMS of localization precision or localization bias, which was calculated from different emitters, as the final localization precision or localization bias.

## 3. Results

### 3.1 Characterization of sCMOS noise

Since sCMOS noises vary from pixel to pixel, it is necessary to measure these noises in every pixels, and thus the characterization of read noise, offset FPN, gain FPN in conventional cameras should be replaced by the characterization of read noise map, offset map, photon response map, respectively. Here, we use relative offset map instead of offset map to compensate the mean offset value difference among cameras. We use photon response map to replace gain map and/or QE map, so that the calculation can be easier. Note that this treatment is accurate enough for photon signal calculation but would lead to a small bias in electron signal calculation [15]. We further analyze these noise maps by: 1) calculate the SD of relative offset map and photon response map to obtain DSNU and PRNU, respectively; 2) calculate the RMS of read noise map to obtain the amplitude of global read noise of the entire camera; 3) calculate the ratio of the SD to the RMS of the read noise to present RNNU.

We characterized two popular back-illuminated sCMOS cameras, including a Prime 95B (SN: A18A203022, Photometrics, USA) and a Dhyana 95 (SN: KBS4951703002, Tucsen Photonics, China), and a representative iXon Ultra 897 EMCCD camera (SN: X-4652, Andor, England). 5000 dark frames were used to calculate the read noise map and relative offset map, and 6 groups of bright images with 1000 frames in each group were used to calculate the photon response map. For these two sCMOS cameras, stripped patterns and high dark noise pixels were easily found in the enlarged noise maps. Here we show only the results from the Dhyana 95 (see Fig. 1a). We further found the camera noises in the iXon 897 are much smaller than those in the sCMOS cameras: 1) The DSNU of the Dhyana 95, Prime 95B and iXon 897 are 1.31 e-, 0.52 e-, 0.07 e-, respectively; 2) The PRNU of the Dhyana 95, Prime 95B and iXon 897 are 1.02 %, 0.65 %, 0.32 %, respectively; 3) the RNNU is ∼ 24% for both sCMOS cameras and ∼2% for the iXon Ultra 897, while the RMS of read noise is ∼2 e-in both sCMOS cameras and 0.43 e-in the iXon Ultra 897 (Fig.1b). Additionally, as compared with the Dhyana 95, we found a much shorter trail in the read noise probability distribution function (PDF) of the Prime 95B. It is probably because some defect pixels (i.e. high read noise pixels) had been corrected by the default defect pixel correction in the Prime 95B.

**Fig. 1.**
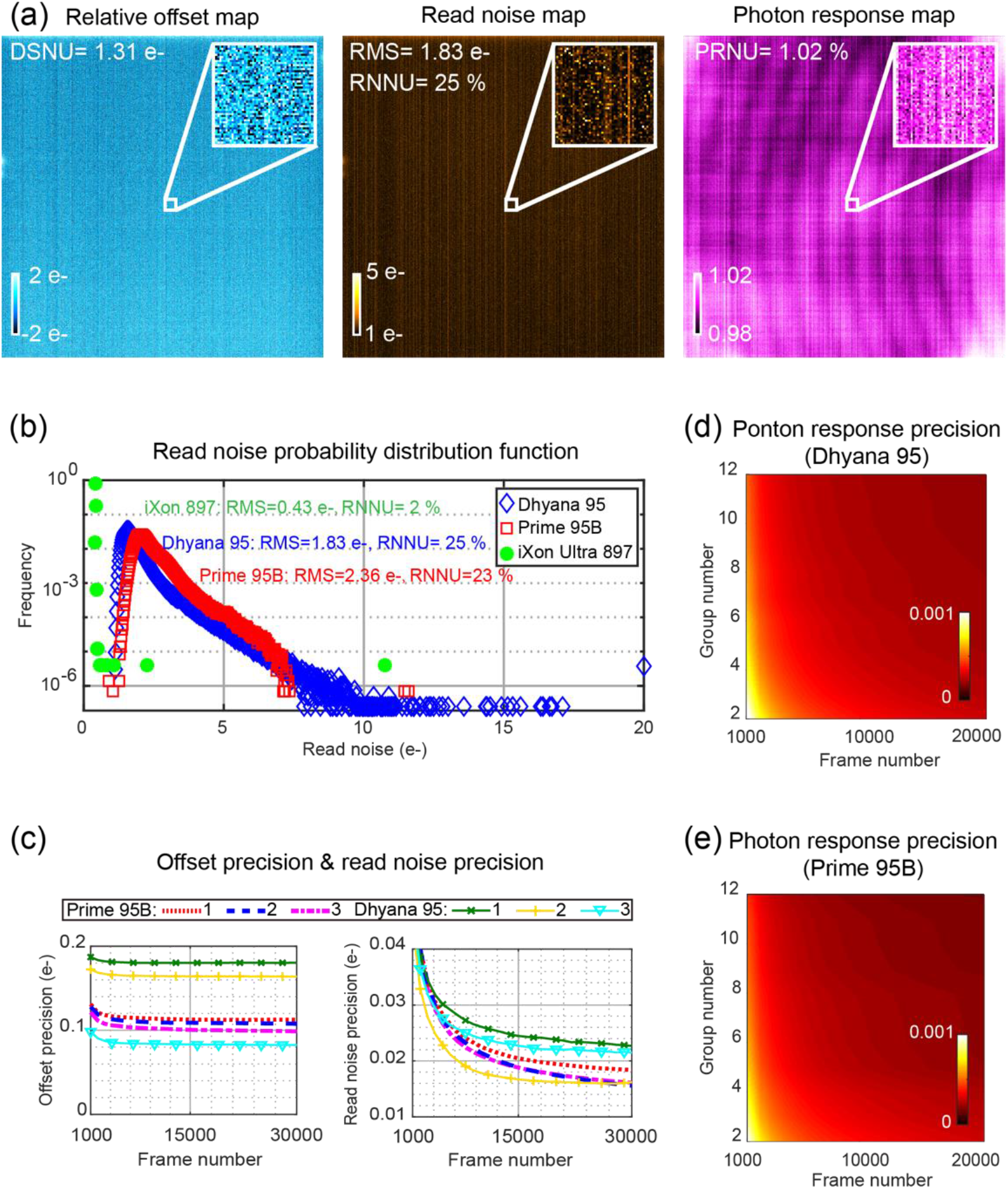
Camera noise characterization. (a) The relative offset map, read noise map, and photon response map of the Dhyana 95. (b) The read noise probability distribution function (PDF) for the three cameras. The dependence of the measurement precision of (c) the relative offset map, read noise map, and (d, e) the photon response map on frame number and group number. The entire images in (a) are 2000 × 2000 pixels and the enlarged images in the top-right corners of (a) are 50 × 50 pixels. Note that the read noise PDF of the iXon 897 has a sharpen distribution. The exposure time was 20 ms.

The dependence of the noise map measurement precision on frame number and group number was characterized in an area of 128×128 pixels, which was cropped from nearly the center of the camera sensor. To measure the precision of the read noise map and relative offset map, six groups of dark frames captured at different days were randomly divided into three groups of datasets. These datasets were further analyzed to present three independent measurements showed in Fig. 1c. Result shows that the Prime 95B has a better repeatability than the Dhyana 95, indicating a better dark noise control in the former. To measure the precision of the photon response map, two datasets were measured at different days for both the Dhyana 95 and the Prime 95B (Fig. 1d, e). Result shows the photon response map precision are nearly the same for both sCMOS cameras. We found a total of 5000 dark frames are sufficient to provide a precision of < 0.2 e-in the relative offset map and ∼ 0.03 e-in the read noise map, and 6 groups of 1000 bright frames are enough to provide a precision of 0.05% in the photon response map. Note we measured the noise map precision with 20 ms exposure time and the precision will change with exposure time.

We investigated the defect pixels in the Dhyana 95. We considered a pixel as defect pixel if the amplitude of any camera noises in that pixel exceeds 10% of the maximum signal range in the working camera mode. Only the center 1800 × 1800 pixels were used for this characterization, because the performace of the pixels in camera edge is genarally worse than the pixels in camera center for long exposure time. We found the number of low gain pixels or high read noise pixels is much smaller than that of high relative offset pixels. Actually we found 18 low gain pixels and 9 high read noise pixels, but 7093 high relative offset pixels. We also investigated the dependence of relative offset and read noise on exposure time (Fig. 2b). Since dark current increases with exposure time, read noise and offset will also increase accordingly, but the increase of offset will be more obvious. As seen in Fig. 2b, when the exposure time in pixel 1 increases from 1 ms to 10 s, the relative offset increases from ∼ 0 e- to ∼ 480 e-, while the read noise increases from ∼ 2 e- to ∼ 18 e-. In some pixels, the offset even reaches the maximum signal range, leaving no capability for these pixels to response to any incident photons. These findings indicate sCMOS users should pay more attention to offset FPN correction rather than read noise correction when long exposure time is necessary.

**Fig. 2.**
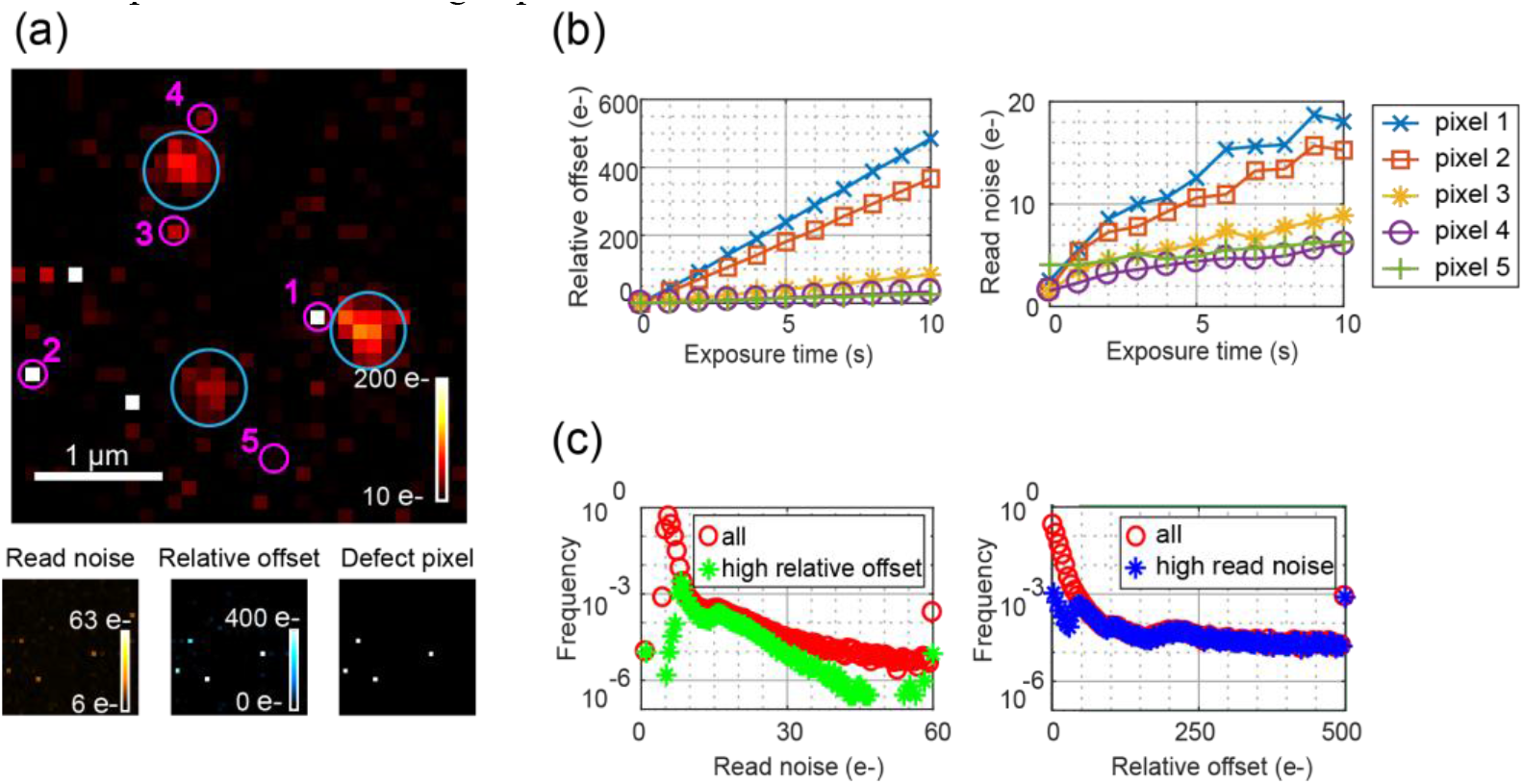
High noise pixel characterization in the Dhyana 95. (a) A fluorescence image of beads (in blue circles). The read noise map and the relative offset map of the same imaging area and the same exposure time are also shown below. The defect pixel map shows the pixels with > 200 e-noise. (b) The dependence of relative offset and read noise on exposure time. Pixel 1-5 were marked out in (a). The data points in (b) were calculated from 100 dark frames. (c) The PDF of read noise (left) and relative offset (right). The statistics in (c) were from the top 1% pixels with highest read noise (high read noise) or offset (high relative offset), or all of the 1800 × 1800 pixels (all). The read noise and the relative offset in (a, c) were calculated from 1000 dark frames. Note the zero read noise pixels in (c) are probably defect pixels whose values are close to the maximum signal range and never change. The exposure time in (a, c) was 10 s.

It is interesting to further investigate whether high read noise pixels overlap with high offset pixels, and whether the indentity of high dark noise pixels changes with exposure time. Using PDF, we identified four groups of high dark noise pixels (top 1% of the high read noise pixels or high offset pixels, with 20 ms or 10 s exposure time), and compared the overlap pixels of any two groups from these four groups. The overlap rate between high read noise pixels and high offset pixels is 57% when the exposure time is 10 s (Fig. 2c), and becomes lower (10%) when the exposure time is 20 ms. When the exposure time changes from 20 ms to 10 s, the location of top 1% high read noise pixels or high relative offset pixels also change (overlap rate < 10%), meaning the identity of high dark noise pixels should be characterized separately for short exposure time and long exposure time.

### 3.2 The impact of different sCMOS noises on conventional microscopy and SMLM

We analyzed the impact of sCMOS noises using simulated images on two imaging scenarios: conventional microscopy and SMLM (Fig. 3). The impact of sCMOS noises on conventional microscopy is directly assessed by the imaging quality of raw image frames, including peak signal-noise-ratio (PSNR), structural similarity (SSIM), and the number of “outlier pixel” (N_OP_). The impact of sCMOS noises on SMLM is assessed by the localization results calculated from raw image frames, including localization precision and localization bias.

**Fig. 3.**
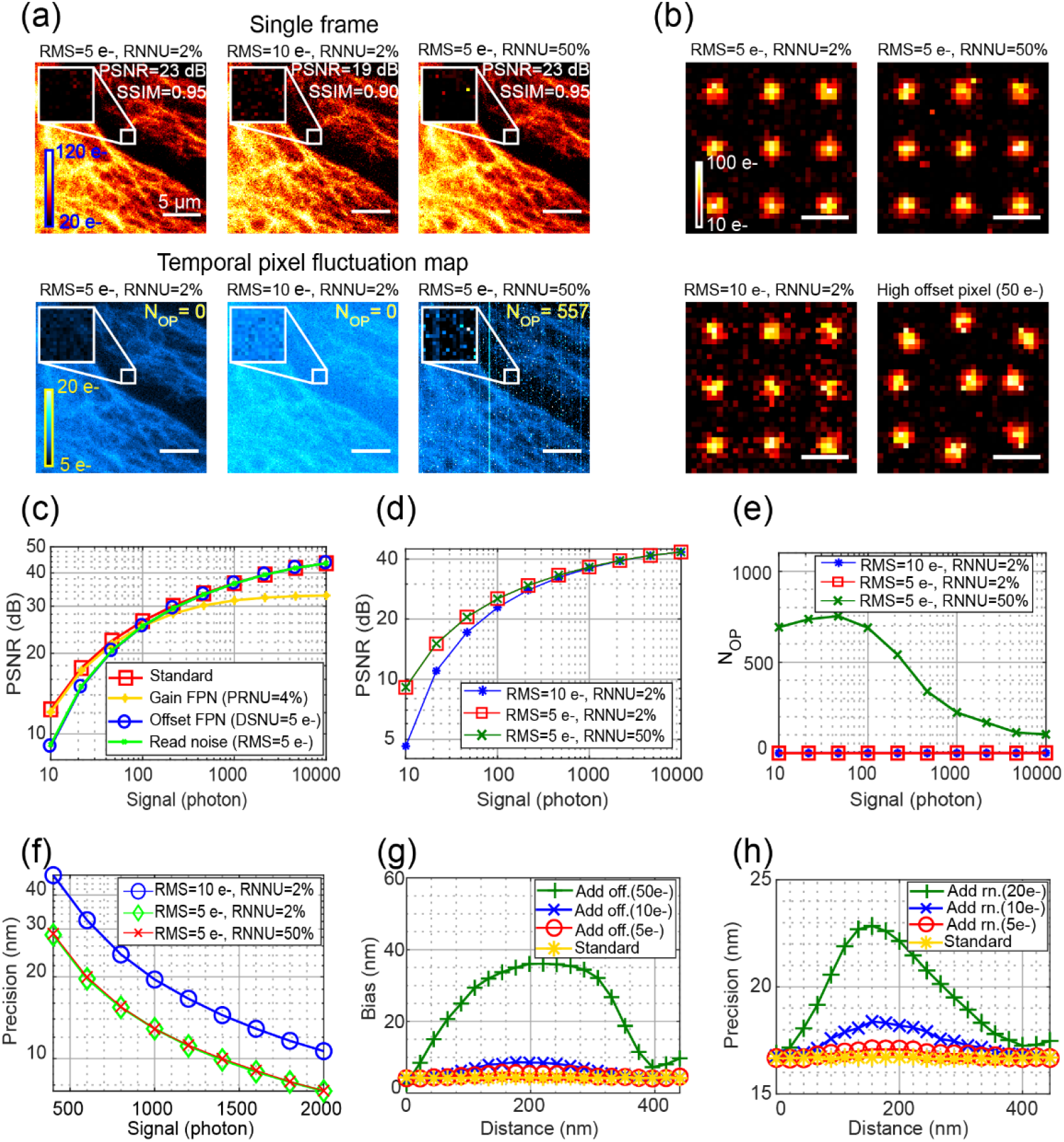
The impact of camera noises on conventional microscopy and SMLM. (a) Three simulated conventional microscopy images with different read noise maps. The corresponding temporal pixel fluctuation maps are shown in the bottom. (b) Four simulated single molecule images. (c) The impact of different camera noises on PSNR. The dependence of (d) PSNR and (e) the number of outlier pixels (N_OP_) on different signal levels for images with different read noise maps. (f) The impact of read noise on localization precision. The dependence of (g) localization bias or (h) localization precision on the distance between high noise pixel and the emitter center. A normal pixel was replaced by a high offset pixel (Add off.) or a high read noise pixel (Add rn.) in the standard noise maps around each emitter in (g) or (h). The simulated noise maps in (a-h) were the noise maps of the Dhyana 95 (that is, the standard noise maps, DSNU = 1.31 e-, PRNU = 1.02 %, RMS of read noise = 1.83 e-, RNNU = 25%) or changing a noise map from the standard noise maps. The changed noise map was shown in the legend. The simulated background in (a-h) is 10 photons/pixel. The PSNR and SSIM and the temporal pixel fluctuation maps in (a, c-e) were calculated from 100 frames for each group. The signal level in (a) is ∼63 photons/pixel. The signal of the emitters in (b, f-h) is 500 photons/emitter. Pixel size in (a-b): 110 nm. Scale bar in (b): 1 μm.

For conventional microscopy images, we used the measured noise maps (including read noise map, relative offset map, and photon response map) of the Dhyana 95 as standard noise maps, and enhanced the noise amplitude in these maps to obtain high noise maps. We generated high noise images with a high level of camera noise (including global read noise, offset FPN, or gain FPN), using a ground-truth image, the standard noise maps and the high noise maps. Details can be found in Section 2.5. We calculated PSNR and SSIM using the simulated images with standard and high noise maps, and found offset FPN and read noise decrease PSNR when the signal is low (< 100 photons), and gain FPN decreases PSNR when the signal is relatively high (> 100 photons) (Fig. 3c). Similar results were found for the SSIM assessment. Note that, for both the PSNR and SSIM assessment, the common-used reference image is the averaged image without FPN correction. But this image could not be used to assess the impact of FPN, because averaging images could only eliminate temporal random noise rather than FPN.

We compared the impact of global read noise and RNNU on the image quality of conventional microscopy. To visualize the impact more easily, we simulated images with several times higher RMS of the read noise maps, as compared with the standard read noise map. We studied three groups of simulated images with different read noise maps and used the relative low read noise group (RMS = 5 e-, RNNU= 2%) as the reference. We found the PSNR of the high global read noise group (RMS = 10 e-) is the lowest when the signal is moderate or low (< 1000 photons), and the PSNR of the high RNNU group (RNNU = 50%) is close to that of the reference group in the full signal range under studied (Fig. 3d). However, only the high RNNU group has outlier pixels (Fig. 3e). These findings indicate: 1) global read noise decreases the image quality of the whole image, while RNNU raises a larger temporal fluctuation in some high read noise pixels, including but not limited to the outlier pixels; 2) PSNR is not suitable to assess the impact of RNNU on image quality. Because FPN would not change with time, it adds only a fixed bias to the digital value of a pixel, and thus would not increase N_OP_. Therefore, for the camera noises under studied, N_OP_ only increases with RNNU.

For SMLM, FPN was previously found to increase localization bias, while read noise would degrade localization precision [11–13]. Here, we compared the impact of RNNU and global read noise on localization precision by simulating three groups of single molecule images with three different read noise maps. We found the localization precision of the high global read noise group (RMS = 10 e-) is the lowest, and the localization precision of the high RNNU group (RNNU = 50%) is the same as the reference group (RMS = 5 e-, RNNU = 2%) (Fig. 3f). This is probably because the read noise map with high RNNU has not only more relative high read noise pixels but also more low read noise pixels, and their impact on localization precision is counteracted from a statistical point of view.

We also studied the impact of isolated high dark noise pixels on SMLM. We simulated single molecule images with high noise maps by replacing a normal pixel with a high read noise pixel or a high offset pixel, with a fixed amplitude and varied distance in the standard noise maps around each emitter. We found the impact of the isolated high noise pixel changes with the distance between the emitter center and the position of the high noise pixel (Fig. 3g-h). A high offset pixel with 10 e-relative offset could lead to >1 nm localization bias, while a high read noise pixel with 10 e-read noise could lead to ∼ 10% degradation in localization precision. For the Dhyana 95 with 20 ms exposure time, there are only ∼ 60 pixels with read noise > 10 e- and ∼ 20 pixels with relative offset > 10 e-in an area of 2000 × 2000 pixels, meaning the high dark noise pixels bring a negligible degradation to the performance of SMLM. However, for some unlikely cases where the exposure time increases to several seconds, the number of high dark noise pixels may increase significantly (see Fig. 2b-c) and notable degradation for SMLM may be observed.

### 3.3 FPN correction

We captured 5000 dark images to calculate a DSNU map, and 5000 bright images at ∼950 e-/pixel to calculate a PRNU map, and used the statistic parameters (including DSNU, PRNU, and local sensitivity variation (SV)) to assess the performance of FPN correction. For the two sCMOS cameras, we directly used the measured relative offset maps and photon response maps to perform FPN correction. After correction, the DSNU decreases from 1.31 e-to 0.16 e-for the Dhyana 95 and from 0.52 e-to 0.15 e-for the Prime 95B (Fig. 4a), respectively. For both sCMOS cameras, the DSNU after correction is already close to quantizing noise (that is, 0.14 e-for the Dhyana 95 and 0.17 e-for the Prime 95B) [19], which is the minimum noise for digital equipment. The DSNU for the iXon 897 was measured to be 0.07 e-before correction and < 0.01 e-after correction, when the EMgain is 100. Note the corrected DSNU of the iXon 897 is mainly limited by quantizing noise (∼ 0.03 e-). For gain FPN, the corrected PRNU and SV of both sCMOS cameras decrease to < 0.3% and < 0.2%, respectively (Fig. 4 b), which are comparable to those of the iXon 897 (before correction: PRNU = 0.32%, SV=0.22%; after correction: PRNU = 0.12%, SV = 0.07%). In addition, all of the characterization parameters (DSNU, PRNU, SV) of the uncorrected Prime 95B are better than those of the Dhyana 95, and the uncorrected SV of the Prime 95B is even close to that of the iXon 897. These findings suggest the FPN of the Prime 95B had been partially corrected in the factory, and could be further corrected by end-users if necessary. Therefore, we verified that FPN could be corrected to a negligible level using a proper method.

**Fig. 4.**
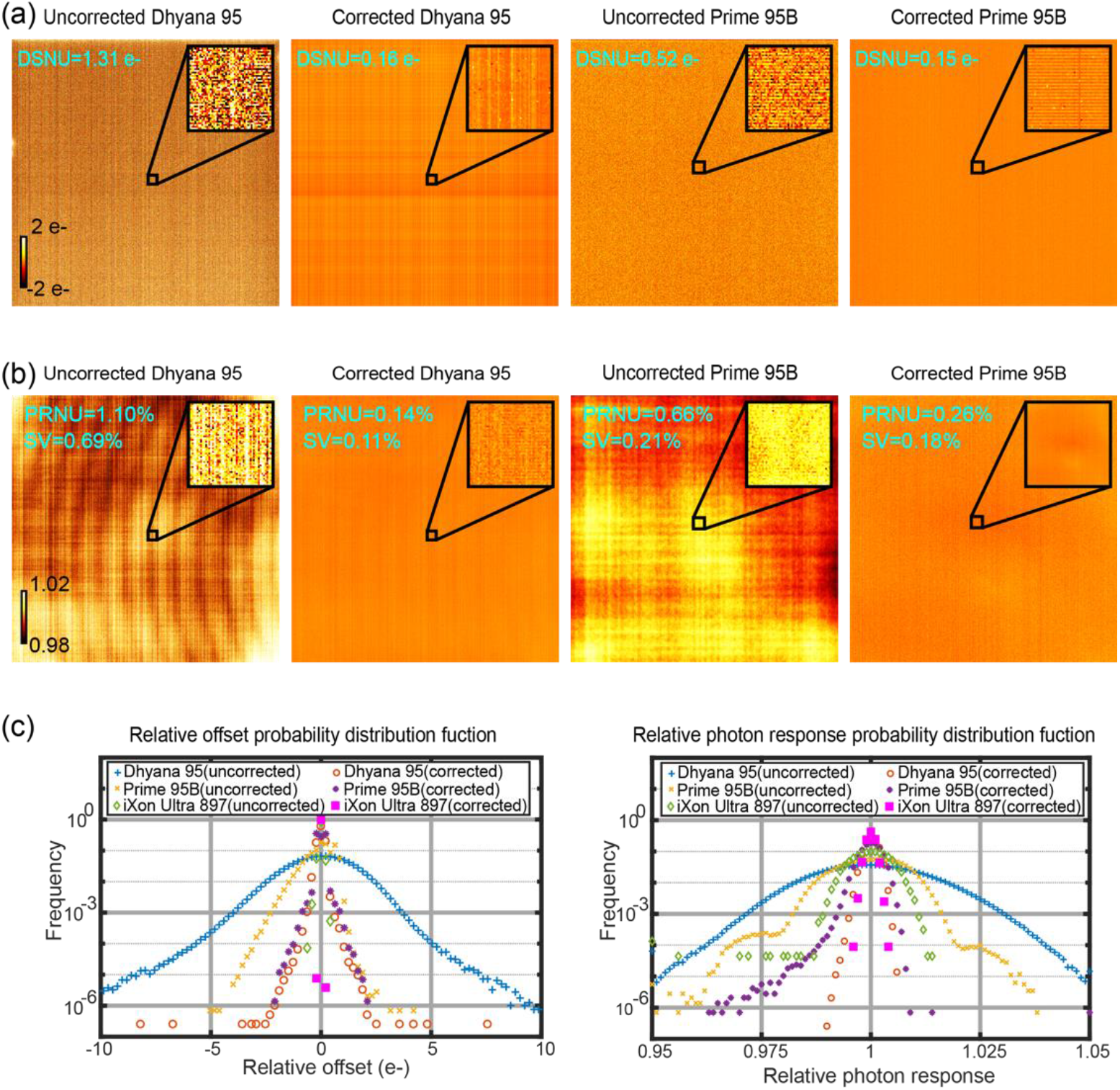
FPN correction for the Dhyana 95 and the Prime 95B. (a) DSNU map. (b) PRNU map. (c) The corresponding PDF of the relative offset (DSNU map) and the relative photon response (PRNU map). The DSNU and PRNU results in (a-c) were averaged from 5000 images. The whole images in (a-b) are 2000 × 2000 pixels for the Dhyana 95, and 1200 × 1200 pixels for the Prime 95B. The exposure time was 20 ms.

We further analyzed the robustness of FPN correction (DSNU and PRNU) using the Dhyana 95. Offset FPN correction may not work well when the experimental images are not captured with the same exposure time as the relative offset map, because the offset value of a pixel increases with exposure time. For example, when a relative offset map with 1 s exposure time was used to correct the DSNU map with 20 ms exposure time, the DSNU was found to increase from 1.31 e-to 3.46 e-.

For gain FPN correction, different experimental conditions were analyzed (Table 1). Typically, we performed gain FPN correction using photon response maps measured under the following conditions: 600 nm illumination wavelength, approximately 150 ∼ 1700 e-/pixel signal intensity, 20 ms exposure time. We found the performance of gain FPN correction would degrade when the raw images are not taken with the same experimental conditions used for the photon response map measurement. Specifically, we found the photon response map measured under the same illumination wavelength could decrease the PRNU from > 1% to < 0.15%, but a photon response map measured under a different illumination wavelength (600 nm) could even increase the PRNU (measured under 850 nm) from 1.57% to 1.64%. However, the photon response map measured under a mismatched illumination wavelength could still be effective to correct SV from ∼ 0.69% to ∼ 0.12%, which is close to the case using matched experimental conditions. We also found the photon response map measured under mismatched intensities could partly correct the PRNU and SV, but the photon response map measured under matched intensities provides much better correction. These findings prove that, to obtain an optimal FPN correction, both the relative offset map and photon response map should be measured under the same experiment conditions.

**Table 1.** Gain FPN correction using different photon response maps^*a*^.

| Parameters | Correction Type<sup>b</sup> | Standard | 850 nm<sup>c</sup> | 400 nm<sup>c</sup> | Intensity<sup>d</sup> | Exposure time<sup>e</sup> |
| --- | --- | --- | --- | --- | --- | --- |
| PRNU | No correction | 1.10% | 1.57% | 1.07% | 1.74% | 1.16% |
|  | Mismatched | - | 1.64% | 0.35% | 1.10% | 0.28% |
|  | Matched | 0.14% | 0.13% | 0.15% | 0.27% | 0.16% |
| SV | No correction | 0.69% | 0.68% | 0.70% | 1.35% | 0.74% |
|  | Mismatched | - | 0.12% | 0.12% | 0.95% | 0.20% |
|  | Matched | 0.11% | 0.11% | 0.11% | 0.23% | 0.13% |

<sup>a</sup>The photon response map measured under typical conditions (600 nm illumination wavelength, appropriate 150 e-~1700 e-/pixel intensity level, 20 ms exposure time) was used as the standard map. The PRNU map was measured under the typical conditions (standard) or customized conditions where one parameter in the typical conditions was changed and others remained the same. <sup>b</sup>The PRNU map was corrected by the standard map (mismatched) or the photon response map measured under the same conditions as the PRNU map (matched). <sup>c</sup>PRNU map measured under 400 nm or 850 nm illumination. <sup>d</sup>PRNU map measured under an illumination intensity of ~ 105 e-/pixel, and the matched
photon response map was measured under an illumination intensity of appropriate 30 ~ 200 e-/pixel. PRNU map measured under 1 s exposure time.

It is worthy to note that, during the past two years we repeated the FPN correction with the Dhyana 95 several times, and found the offset map changes only slightly, while the photon response map is kept stable. When the relative offset map and the photon response map were measured several months before used, the DSNU after correction changed from 0.2 e-to ∼ 0.5 e-, while the SV after correction was still nearly the same (from 0.11% to 0.13%). So the relative offset map should be checked and updated regularly. Besides, because PRNU is sensitive to the illumination angle that may changes after a long period (for example, several months), the corrected PRNU of the Dhyana 95 may deteriorate from 0.15% to ∼ 0.30%, depending on the illumination angle repeatability.

### 3.4 Defect pixel correction

We corrected the defect pixels in the Dhyana 95 with a 3 × 3 average filter, and compared the results with those from FPN corretion (Table 2). We assessed the performance of these corrections under different exposure time: 20 ms or 1 s or 10 s for DSNU, and 20 ms for PRNU. The DSNU map with 1 s exposure time was averaged from 5000 dark frames, and the DSNU map with 10 s exposure time was averaged from 1000 dark frames. Results show the improvement in DSNU and PRNU is negligible when the exposure time is 20 ms. With a longer exposure (1 s or 10 s), defect pixel corretion improves DSNU, but FPN correction improves DSNU more significantly. Moreover, a combination of FPN correction and defect pixel correction brings a better improvement in DSNU than FPN or defect pixel correction itself. Note the FPN correction needs to be performed with noise maps under matched experimental conditions, while the defect pixel correction doesn’t require a tight experimental control. This brings special cautions for the correction under long exposure time. Actually, using a relative offset map with 10 s exposure time to correct the DSNU map with 1 s exposure time will increase the DSNU from 2.94 e-to 24.24 e-. However, if we use the relative offset map with 10 s exposure time to determine the defect pixels and then perform defect pixel correction for the DSNU map with 1 s exposure time, the DSNU would decrease from 2.94 e-to 1.46 e-. This means defect pixel correction is easier to perform than FPN correction for experiments with varied exposure time.

**Table 2.** Comparing defect pixel correction with FPN correction in the Dhyana 95.

| Parameters | Exposure time | Without correction | Defect pixel correction | FPN correction | Combined correction<sup>a</sup> |
| --- | --- | --- | --- | --- | --- |
| PRNU | 20 ms | 1.04% | 1.04% | 0.14% | 0.14 % |
| DSNU | 20 ms | 1.31 e- | 1.30 e- | 0.16 e- | 0.16 e- |
| DSNU | 10 s | 27.92 e- | 9.34 e- | 3.94 e- | 3.53 e- |
| DSNU | 1 s | 2.94 e- | 1.46 e- | 0.80 e- | 0.76 e- |

<sup>a</sup>The combined correction executes firstly FPN correction, then defect pixel correction.

### 3.5 Scenario-specific noise correction for conventional microscopy

We compared the performance of NCS [16] (an sCMOS-specific de-noising algorithm) with PURE [27] (a normal de-noising algorithm) by correcting the same group of experimental images: 100 frames of fluorescent microtubules in U-2 OS cells. We found both algorithms could increase the PSNR and SSIM, but PURE is more effective. Compared with NCS, PURE brings more smooth structures in the corrected images (Fig. 5a) and a lower temporal fluctuation (Fig. 5b). However, as seen in Fig. 5b, the outlier pixels can be well-corrected by NCS, but not PURE. This means that, for conventional microscopy images, NCS is effective in minimizing the impact of RNNU.

**Fig. 5.**
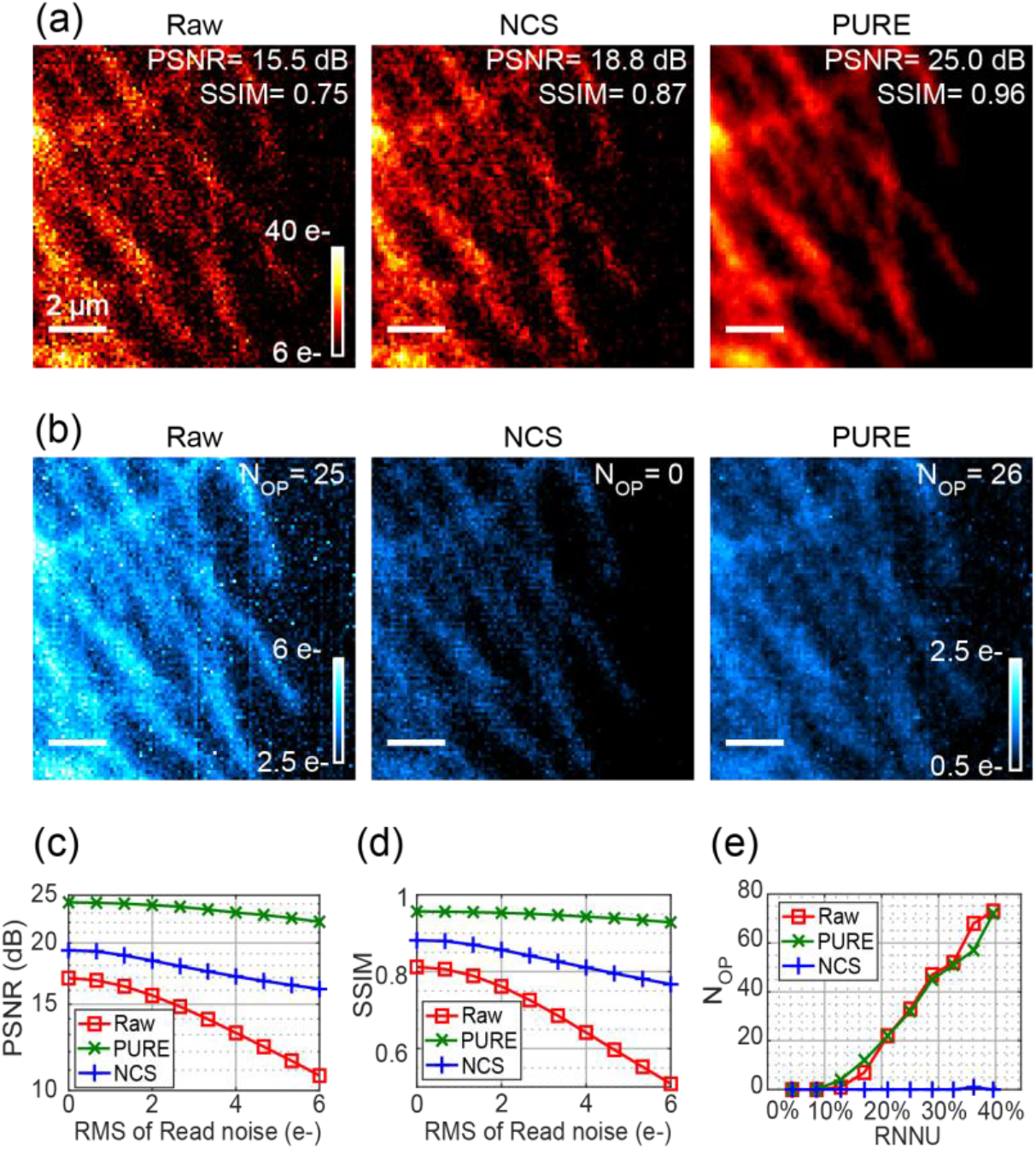
Noise correction for conventional microscopy. (a) A raw image and the corresponding corrected images using an sCMOS-specific de-noising algorithm (NCS) and a normal de-noising algorithm (PURE). (b) Three corresponding temporal pixel fluctuation maps. The individual bright pixels in (b) may be outlier pixels. The dependence of (c) PSNR and (d) SSIM on the RMS of read noise. (e) The dependence of N_OP_ on RNNU. The PSNR, SSIM and temporal pixel fluctuation maps in (b-e) were calculated from 100 frames for each group. In (b), the left and the middle maps share the same color bar marked in the left, and the right map has a different color bar. The pixel size in (a-b) was 110 nm.

We further compared the performance of these two de-noising algorithms using simulated images, which are based on the same group of experimental images, but with different noise maps. We simulated images with different RMS of read noise, and found both the PSNR and SSIM of the corrected images decrease with the increase of RMS of the read noise for both de-noising algorithms (Fig. 5c-d). This means the image quality of the corrected images is negatively correlated with the global read noise. To further improve the image quality, new algorithms or better circuit design should be developed to minimize global read noise. On the other hand, for both de-noising algorithms, we found the PSNR or SSIM improves even when the RMS of read noise is ∼ 0 (Fig. 5c-d), indicating that these two algorithms minimize not only global read noise but also shot noise. Additionally, using simulated images with different RNNU, we verified NCS could always correct the outlier pixels (Fig. 5e), confirming the previous finding that NCS could minimize the impact of RNNU on conventional microscopy images [31].

We performed FPN correction and defect pixel correction on the same group of experimental images. After FPN correction, the PSNR increased from 15.5 dB to 15.8 dB, the SSIM increased from 0.75 to 0.76, and N_OP_ was not changed as expected (Data not shown in Fig. 5). Although in the FPN correction section we confirmed FPN could be corrected to a negligible level, the increase of PSNR and SSIM from FPN correction is not as remarkable as those from NCS or PURE (see Fig. 5a). That is because temporal noises (including read noise and shot noise) are the dominant noises under this signal intensity [5]. Note that NCS performs FPN correction at the first step. On the other hand, after defect pixel correction, the PSNR, SSIM, and N_OP_ became 16.3 dB, 0.79, and 25 (Data not shown in Fig. 5), respectively. The N_OP_ didn’t change because the defect pixels determined under long exposure time may not overlap with the relative high read noise pixels with short (20 ms) exposure time, and thus was not corrected. Compared with the NCS and PURE, the FPN correction and defect pixel correction keep more raw data, at the expense of having less impact on imaging quality. Taking these results together, we conclude that global read noise would be the limit for noise correction, and the impact of RNNU on conventional microscopy images could be well corrected by NCS.

### 3.6 Scenario-specific noise correction for SMLM

We analyzed the localization algorithms with the simulated single molecule images mentioned in Section 3.2. We used MLE_sCMOS_ to localize the molecules simulated with different read noise maps and compared the results with those from MLE_normal_ (Fig. 6a). We found MLE_sCMOS_ improves ∼ 10% in localization precision in the high RNNU group, but does not change the localization precision in the low RNNU group. Because there are more relative high read noise pixels in the high RNNU group, the degradation of high read noise pixels on localization precision could be compensated by MLE_sCMOS_. For the low RNNU group, there are no relative high read noise pixels, and thus the localization precision doesn’t benefit from MLE_sCMOS_. These results indicate MLE_sCMOS_ could correct the impact of RNNU but not global read noise on localization precision.

**Fig. 6.**
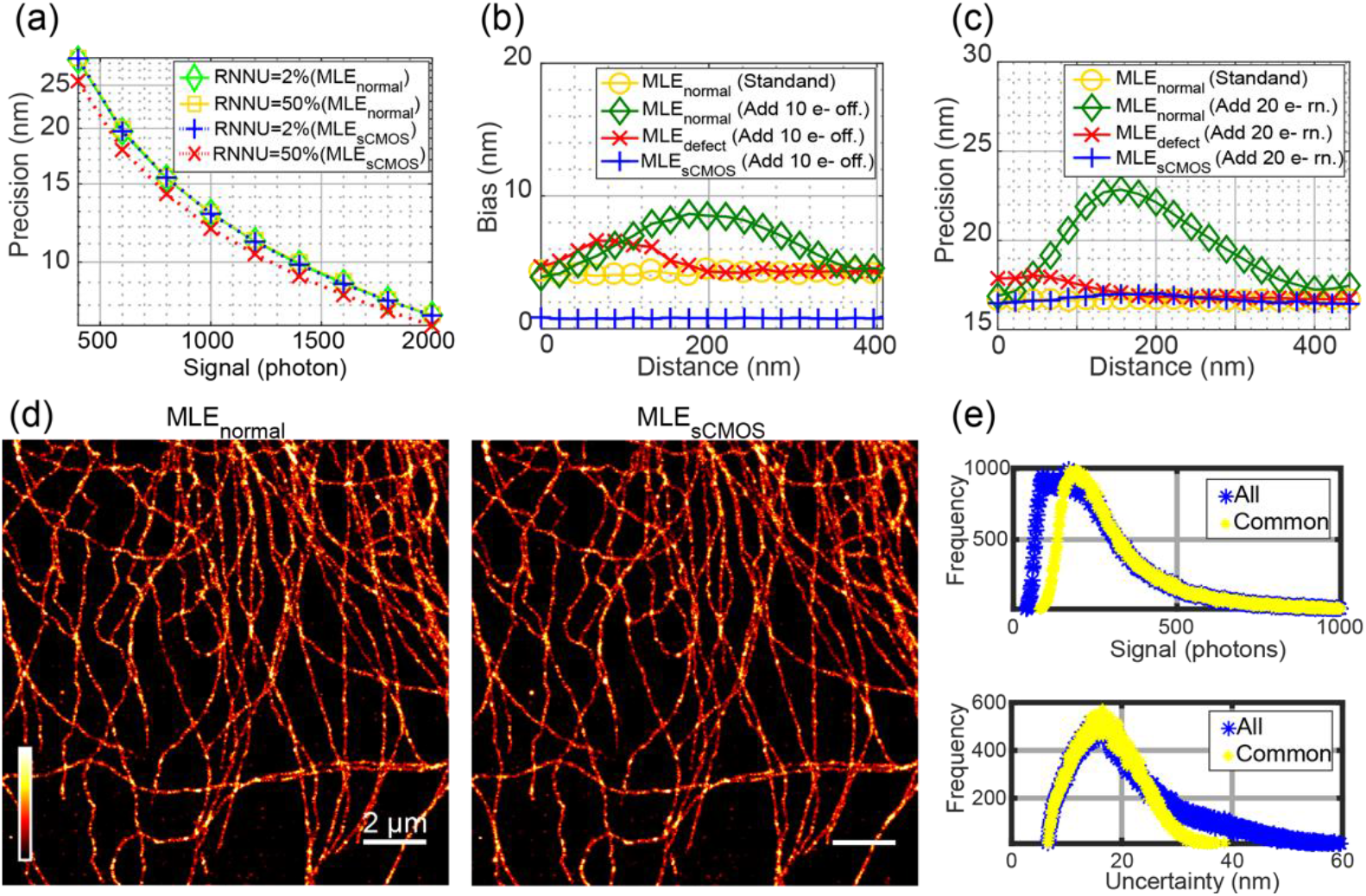
The performance of three localization algorithms (MLE_normal_, MLE_sCMOS_, and MLE_defect_) for SMLM. (a) Localization precision of MLE_sCMOS_ and MLE_normal_ for simulated images with different RNNU. (b) Localization bias and (c) localization precision of the three algorithms for simulated images with isolated high noise pixels: 10 e-relative offset pixels in (b), and 20 e-read noise pixels in (c). (d) Rendered super-resolution images. (e) Localization statistics of all molecules localized by MLE_sCMOS_ (All) and the molecules localized by both MLE_sCMOS_ and MLE_normal_ (Common). All of the localization statistics data in (e) were originated from MLE_sCMOS_ calculation results. The signal in (a-c) was 500 photons/emitter. In (d), the image size was 128× 128 pixels, and the pixel size was 110 nm.

We examined the performance of MLE_sCMOS_ on an extreme case, where the emission pattern from single molecules is contaminated by an isolated high noise pixel. We also evaluated the performance of a modified MLE_normal_ (called MLE_defect_), where MLE_normal_ is used to localize the raw images after defect pixel correction. We used the three algorithms (MLE_normal_, MLE_sCMOS_ and MLE_defect_) to localize two selected groups of images (10 e-relative offset group, and 20 e-read noise group) in Fig. 3g-h, and then compared their localization results with the standard group in Fig. 3g-h. We found MLE_sCMOS_ decreases the localization bias from > 8 nm to ∼ 1 nm (Fig. 6b), which is obviously lower than that of the standard group. That is because MLE_sCMOS_ performs FPN correction before localization, which corrects the fixed bias from not only the isolated high offset pixel but also every pixel. We also found MLE_sCMOS_ could improve localization precision from > 23 nm to ∼ 16 nm (Fig. 6c), which is slightly worse than the localization precision of the standard group. That means MLE_sCMOS_ is effective to correct the impact of isolated high read noise pixel on localization precision. Since the high read noise pixels in sCMOS cameras are usually random distributed, we conclude MLE_sCMOS_ could minimize the impact of high read noise pixels on SMLM.

On the other hand, MLE_defect_ could reduce the impact of isolated high noise pixels on the localization bias and localization precision, especially when the isolated high noise pixel is 100 nm (or more) away from the emitter center (Fig. 6b-c). However, as compared with the standard group, MLE_defect_ was observed to have an increase of ∼ 2 nm in the localization bias and localization precision (Fig. 6b-c). This is probably because the digital value of the isolated high noise pixel was replaced by the mean value of 8 adjacent pixels, which reduces the impact of high noise pixels on SMLM, but results in a distorted emission pattern. That is to say, when normal pixels are considered as defect pixels and are corrected, the localization precision and localization bias would be degraded when a normal MLE algorithm is used.

We compared MLE_sCMOS_ and MLE_normal_ by localizing the same group of SMLM data: 10000 raw image frames captured by the Dhyana 95 with 1 ms exposure time (Fig. 6d). We found MLE_scmos_ could localize more molecules than MLE_normal_: 188890 molecules for MLE_scmos_ and 161464 molecules for MLE_normal_. This is probably because MLE_scmos_ considers noise non-uniformity when image segmentation is performed [11]. We further identified 157559 molecule pairs from both localization algorithms, and compared them with all of the molecules localized by MLE_scmos_ (Fig. 6e). We found most of the molecules localized by MLE_scmos_ but not MLE_normal_ are low signal molecules, which are beneficial for fast SMLM imaging. But this advantage is not enough to raise a notable difference in the global spatial resolution when the number of molecules is high. Actually, the Fourier ring correlation (FRC) [32] of the final super-resolution image is close: 135.6 nm for MLE_normal_, and 137.4 nm for MLE_scmos_.

Taking these results together, it is reasonable to conclude that MLE_sCMOS_ is a good choice for localizing raw images with high noise non-uniformity, because MLE_sCMOS_ could minimize the impact of RNNU but not global read noise on SMLM. This conclusion also agrees with the theoretical analysis of the localization precision in sCMOS camera, where read noise map should be included [11, 15].

## 4. Discussion and conclusion

Camera noise non-uniformity is a major concern for the selection and use of sCMOS cameras. In this paper, we analyzed systematically different kinds of camera noises in two popular back-illuminated sCMOS cameras, and confirmed camera noise non-uniformity (including offset FPN, gain FPN, and RNNU) could be well-corrected by using proper algorithms. We also studied the impact of different noises on conventional microscopy and SMLM, and investigated the usability of FPN correction and defect pixel correction performed by camera manufactories.

We found the commonly-used parameters (including PSNR and SSIM for convectional microscopy, localization precision and FRC for SMLM) are insensitive for assessing the noise non-uniformity, and thus new methods or parameters should be developed to characterize the impact of the noise non-uniformity on imaging quality. We suggest to study the regions around high noise pixels separately when discussing the impact of sCMOS noise non-uniformity on imaging quality.

Both defect pixel correction and FPN correction are regularly used by camera manufactories to improve the image quality of sCMOS cameras. However, their usability should be considered carefully. For defect pixel correction, manufactory usually determines defect pixels using images with long exposure time, because the defect pixels are more obvious under long exposure time. However, because dark noise increases with exposure time, some defect pixels identified under long exposure time may be recognized as normal pixels when the exposure time is short. Taking defect pixel correction to these normal pixels results in distorted images.

For FPN correction, because offset FPN may change over time, relative offset map should be measured regularly. The effectiveness of FPN correction depends on whether the noise maps are measured with the same conditions as the experiment, FPN re-correction may be necessary for a specific experiment. Besides, although the number of high noise pixels in the Dhyana 95 changes little during the past two years, we did observe some newly developed high offset pixels in another sCMOS camera that has been used for six years. We recommend sCMOS users to keep tracking the number and locations of high noise pixels, and perform re-correction if necessary.

Although camera noise non-uniformity can be well-corrected, performing the correction may need additional experiments and expertise, and using the correction algorithms could be time-consuming. Therefore it is necessary to determine whether any sCMOS noise correction algorithms should be used in a specific imaging scenario. After considering all the findings in this paper, we present the following suggestions: 1) For most sCMOS users, it is necessary to perform noise non-uniformity correction only when the final images or the post processing algorithms cannot tolerate the isolated high noise pixels (like pixel 1-2 in Fig. 2a), because noise non-uniformity has smaller impact on image quality than shot noise and global read noise for normal pixels. 2) FPN correction is necessary for some applications that requires time-domain averaging to improve the experiment precision, because in this case shot noise and global read noise have been minimized. A typical example is ultra-high resolution imaging of nuclear pore complex scaffold via particle averaging, where the localization precision is expected to be better than 1 nm [33]. 3) For experiments with long exposure time, dark FPN correction is always recommended, except for varied exposure time in continuous frames.

Finally, it is worthy to point out that, after all possible corrections, the camera noise non-uniformity is no longer the major problem for applying sCMOS cameras in various application fields. To further improve the imaging performance of current commercial sCMOS cameras, camera manufactories and end-users can take more efforts to minimize the RMS of read noise in sCMOS cameras. Recently, Hamamatsu Photonics took a desirable step towards this direction, and released a low-noise back-illuminated sCMOS camera called ORCA-Fusion BT, where the RMS of read noise is 0.7 e-for low read out speed mode. It would be interesting to see what kind of new applications would benefit from this technology advance.

## References

1. X. Michalet, R. A. Colyer, G. Scalia, A. Ingargiola, R. Lin, J. E. Millaud, S. Weiss, O. H. Siegmund, A. S. Tremsin, J. V. Vallerga, A. Cheng, M. Levi, D. Aharoni, K. Arisaka, F. Villa, F. Guerrieri, F. Panzeri, I. Rech, A. Gulinatti, F. Zappa, M. Ghioni, and S. Cova, “Development of new photon-counting detectors for single-molecule fluorescence microscopy,” Philos Trans R Soc Lond B Biol Sci 368, 20120035 (2013).

2. M. Baker, “Faster frames, clearer pictures,” Nature Methods 8, 1005–1009 (2011).

3. Y. J. Wang, L. X. Zhao, Z. Hu, Y. N. Wang, Z. Y. Zhao, L. C. Li, and Z. L. Huang, “Quantitative performance evaluation of a back-illuminated sCMOS camera with 95% QE for super-resolution localization microscopy,” Cytometry Part A 91a, 1175–1183 (2017).

4. B. Moomaw, “Camera technologies for low light imaging: overview and relative advantages,” Methods in Cell Biology 114, 243–283 (2013).

5. L. Li, M. Li, Z. Zhang, and Z.-L. Huang, “Assessing low-light cameras with photon transfer curve method,” Journal of Innovative Optical Health Sciences 09, 1630008 (2016).

6. J. Enderlein, I. Gregor, Z. K. Gryczynski, R. Erdmann, F. Koberling, S. Watanabe, T. Takahashi, and K. Bennett, “Quantitative evaluation of the accuracy and variance of individual pixels in a scientific CMOS (sCMOS) camera for computational imaging,” Proc. SPIE 10071, 100710Z (2017).

7. W. X. Wang, Z. X. Ling, C. Zhang, Z. Q. Jia, X. Y. Wang, Q. Wu, W. M. Yuan, and S. N. Zhang, “Characterization of a BSI sCMOS for soft X-ray imaging spectroscopy,” Journal of Instrumentation 14, P02025 (2019).

8. Y. G. Soskind, A. Marsh, A. Mullan, M. Barszczewski, and J. T. Cooper, “Characterization of performance of back-illuminated SCMOS cameras versus conventional SCMOS and EMCCD cameras for microscopy applications,” Proc. SPIE 10925, 109251C (2019).

9. Z. L. Huang, H. Y. Zhu, F. Long, H. Q. Ma, L. S. Qin, Y. F. Liu, J. P. Ding, Z. H. Zhang, Q. M. Luo, and S. Q. Zeng, “Localization-based super-resolution microscopy with an sCMOS camera,” Optics Express 19, 19156–19168 (2011).

10. S. Saurabh, S. Maji, and M. P. Bruchez, “Evaluation of sCMOS cameras for detection and localization of single Cy5 molecules,” Optics Express 20, 7338–7349 (2012).

11. F. Huang, T. M. Hartwich, F. E. Rivera-Molina, Y. Lin, W. C. Duim, J. J. Long, P. D. Uchil, J. R. Myers, M. A. Baird, W. Mothes, M. W. Davidson, D. Toomre, and J. Bewersdorf, “Video-rate nanoscopy using sCMOS camera-specific single-molecule localization algorithms,” Nat Methods 10, 653–658 (2013).

12. F. Long, S. Q. Zeng, and Z. L. Huang, “Effects of fixed pattern noise on single molecule localization microscopy,” Phys Chem Chem Phys 16, 21586–21594 (2014).

13. R. Lin, A. H. Clowsley, I. D. Jayasinghe, D. Baddeley, and C. Soeller, “Algorithmic corrections for localization microscopy with sCMOS cameras - characterisation of a computationally efficient localization approach,” Opt Express 25, 11701–11716 (2017).

14. C. R. Copeland, J. Geist, C. D. McGray, V. A. Aksyuk, J. A. Liddle, B. R. Ilic, and S. M. Stavis, “Subnanometer localization accuracy in widefield optical microscopy,” Light-Science & Applications 7, 31 (2018).

15. H. P. Babcock, F. Huang, and C. M. Speer, “Correcting Artifacts in Single Molecule Localization Microscopy Analysis Arising from Pixel Quantum Efficiency Differences in sCMOS Cameras,” Scientific Reports 9, 18058 (2019).

16. S. Liu, M. J. Mlodzianoski, Z. Hu, Y. Ren, K. McElmurry, D. M. Suter, and F. Huang, “sCMOS noise-correction algorithm for microscopy images,” Nat Methods 14, 760–761 (2017).

17. B. Mandracchia, X. Hua, C. Guo, J. Son, T. Urner, and S. Jia, “Fast and accurate sCMOS noise correction for fluorescence microscopy,” Nat Commun 11, 94 (2020).

18. A. El Gamal and H. Eltoukhy, “CMOS image sensors,” Ieee Circuits & Devices 21, 6–20 (2005).

19. J. R. Janesick, Photon Transfer (SPIE Press, 2007).

20. J. Leung, J. Dudas, G. H. Chapman, I. Koren, and Z. Koren, “Quantitative Analysis of In-Field Defects in Image Sensor Arrays,” in 22nd IEEE International Symposium on Defect and Fault-Tolerance in VLSI Systems, (2007), pp. 526–534.

21. A. Jain and R. Gupta, “A Survey on Defect and Noise Detection and Correction Algorithms in Image Sensors,” in 2015 International Conference on Advances in Computer Engineering and Applications, (2015), pp. 754–759.

22. “EMVA1288-3.1” (European Machine Vision Association, 2016), retrieved Nov. 9th, 2020, https://www.emva.org/wp-content/uploads/EMVA1288-3.1a.pdf.

23. T. Zhang, X. Li, J. Li, and Z. Xu, “CMOS Fixed Pattern Noise Removal Based on Low Rank Sparse Variational Method,” Applied Sciences 10, 3694 (2020).

24. J. Dudas, C. Jung, G. H. Chapman, K. B. Zahava, and I. Koren, “Robust detection of defects in imaging arrays,” Proc. SPIE 6059, 60590X (2006).

25. H. Gong, S.-x. Xing, Y. Cai, J. Zhang, L. Sun, J.-P. Chatard, B.-k. Chang, and Y.-s. Qian, “Two-point nonuniformity correction based on LMS,” Proc. SPIE 5640, 78541N (2005).

26. G. H. Chapman, S. Djaja, D. Y. H. Cheung, Y. Audet, I. Koren, and Z. Koren, “A self-correcting active pixel sensor using hardware and software correction,” Ieee Design & Test of Computers 21, 544–551 (2004).

27. F. Luisier, T. Blu, and M. Unser, “Image denoising in mixed Poisson-Gaussian noise,” IEEE Trans Image Process 20, 696–708 (2011).

28. M. Ovesny, P. Krizek, J. Borkovec, Z. K. Svindrych, and G. M. Hagen, “ThunderSTORM: a comprehensive ImageJ plug-in for PALM and STORM data analysis and super-resolution imaging,” Bioinformatics 30, 2389–2390 (2014).

29. Z. Y. Zhao, B. Xin, L. C. Li, and Z. L. Huang, “High-power homogeneous illumination for super-resolution localization microscopy with large field-of-view,” Optics Express 25, 13382–13395 (2017).

30. A. Hore and D. Ziou, “Image Quality Metrics: PSNR vs. SSIM,” in 2010 20th International Conference on Pattern Recognition, (2010), pp. 2366–2369.

31. Y. Li, S. Liu, and F. Huang, “Variance lower bound on fluorescence microscopy image denoising,” bioRxiv (2020).

32. N. Banterle, K. H. Bui, E. A. Lemke, and M. Beck, “Fourier ring correlation as a resolution criterion for superresolution microscopy,” Journal of Structural Biology 183, 363–367 (2013).

33. A. Pertsinidis, Y. X. Zhang, and S. Chu, “Subnanometre single-molecule localization, registration and distance measurements,” Nature 466, 647–651 (2010).

